# Volitional Regulation and Transferable Patterns of Midbrain Oscillations

**DOI:** 10.1101/2024.12.04.626827

**Authors:** Hung-Yun Lu, Yi Zhao, Hannah M. Stealey, Cole R. Barnett, Philippe N. Tobler, Samantha R. Santacruz

## Abstract

Dopaminergic brain areas are crucial for cognition and their dysregulation is linked to neuropsychiatric disorders typically treated with pharmacological interventions. These treatments often have side effects and variable effectiveness, underscoring the need for alternatives. We introduce the first demonstration of neurofeedback using local field potentials (LFP) from the ventral tegmental area (VTA). This approach leverages the real-time temporal resolution of LFP and ability to target deep brain. In our study, non-human primates learned to regulate VTA beta power using a customized normalized metric to stably quantify VTA LFP signal modulation. The subjects demonstrated flexible and specific control with different strategies for specific frequency bands, revealing new insights into the plasticity of VTA neurons contributing to oscillatory activity that is functionally relevant to many aspects of cognition. Excitingly, the subjects showed transferable patterns, a key criterion for clinical applications beyond training settings. This work provides a foundation for neurofeedback-based treatments, which may be a promising alternative to conventional approaches and open new avenues for understanding and managing neuropsychiatric disorders.

**Significance statement:** This study demonstrates, for the first time, that neurofeedback using local field potentials (LFP) from the ventral tegmental area (VTA) is feasible in non-human primates. By leveraging the temporal resolution and ability to target deep brain regions, this approach provides a novel way to modulate brain activity linked to dopamine-related functions. The findings reveal that subjects can flexibly control VTA LFP signals and transfer learned strategies to new settings, offering potential for developing neurofeedback-based treatments. This research opens new avenues for managing neuropsychiatric disorders, presenting an alternative to traditional pharmacological interventions that often have side effects and limited effectiveness. The study highlights the plasticity of VTA neurons and their relevance to cognition and mood regulation.

## Introduction

Dopamine is a neurotransmitter that plays a fundamental role in multiple cognitive functions (Soares et al., 2016; Berke, 2018). Midbrain dopaminergic activity from the ventral tegmental area (VTA), an area critical for encoding reward processing information (Pasquereau et al., 2019; Oberto et al., 2023), facilitates learning (Schultz et al., 1997; Steinberg et al., 2013) and mood regulation (Russo and Nestler, 2013; Nikulina et al., 2014). Dysregulation of dopamine function has been strongly linked to psychiatric conditions like depression (Liu et al., 2018). Conventional treatments for depression, such as pharmacological interventions or neuromodulation (e.g., repetitive transcranial magnetic stimulation), can provide relief from symptoms (Witt et al., 2006) by inhibiting dopamine reuptake (Stahl et al., 2004) or inducing dopamine release (Strafella et al., 2001), respectively. However, side effects can occur (Khawam et al., 2006) and the efficacy of these exogenous treatments can vary unpredictably across patients (Voineskos et al., 2020), reflecting our incomplete understanding of how dopaminergic activities are regulated. Gaining insights into the endogenous regulation of VTA activity can shed light on neural mechanisms implicated in regulation and provide a foundation for potential alternative therapeutic approaches.

Neurofeedback, a brain-computer interface (BCI) paradigm where participants observe and modulate their brain activity in real-time (Sitaram et al., 2017), has emerged as a promising technology for treating neurological and psychiatric disorders and investigating brain functions (Fetz, 2013; Moxon and Foffani, 2015). Neurofeedback facilitates modulation of specific neural features and offers a unique opportunity to discern causal relationships between neural activity and behavior (Sitaram et al., 2017). Previous research has demonstrated successful implementation of neurofeedback using electroencephalography (EEG) (Lansbergen et al., 2011; Zoefel et al., 2011; Enriquez-Geppert et al., 2017), functional magnetic resonance imaging (fMRI) (Sulzer et al., 2013a; Watanabe et al., 2017; Keynan et al., 2018), single- and multi-unit activities (spiking activities) (Fetz, 1969; Wyler and Finch, 1978; Ishikawa et al., 2014), and local field potential (LFP) (Engelhard et al., 2013; Khanna and Carmena, 2017; He et al., 2020). However, fMRI’s delayed hemodynamic response (Hellrung et al., 2018), EEG’s difficulty in capturing deep brain activities (Krishnaswamy et al., 2017), and the decreasing amount of recorded neurons over time with spiking activities (Barrese et al., 2013) pose challenges. In the context of VTA neurofeedback, LFP signals outperform EEG, fMRI, and spiking activities. First, LFP has temporal resolution at millisecond scales that matches the dopamine dynamics (Buzsáki et al., 2012). Second, LFP signals exhibit greater stability over time (Ahmadi et al., 2021) and are less prone to signal deterioration (Andersen et al., 2004). Third, LFP can be precisely targeted using a microelectrode array and they directly reflect population activity within the region, with clear relation to net activation and suppression (Buzsáki et al., 2012).

Here, we study VTA LFP neurofeedback in non-human primates (NHPs). Dopaminergic neurons comprise more than 65% of the neurons in the VTA (Holly and Miczek, 2016), suggesting that VTA LFP activity is mainly driven by dopaminergic neurons. We specifically extracted power features from the beta frequency (20-35 Hz) band because it is crucial in encoding information related to reward and decision-making processes (Pasquereau et al., 2019; Oberto et al., 2023), which are functions that dopamine mediates (Bromberg-Martin et al., 2010). For instance, VTA beta power increases during reward-seeking actions and decreases with punishment risk (Park and Moghaddam, 2017), and is modulated in decision-making tasks (Pasquereau et al., 2019; Oberto et al., 2023), suggesting that VTA beta activity correlates with behaviors pathologically impacted in psychiatric disorders.

## Materials and Methods

### Experimental Design

In this study, we trained two rhesus macaque monkeys (Macaca mulatta, n=2) to volitionally up- and down-regulate their normalized LFP beta power in the VTA region by mapping a function of their LFP signal to control the cursor location on a computer screen positioned in front of them (Fig. 1a). Each experimental session consisted of Control, Main, and Washout blocks with an optional Transfer block that followed a successful Main block (Fig. 1b). The Control and Washout blocks reflected a resting state, while the Main and Transfer blocks were the primary assay blocks of interest where the subjects controlled the cursor through a BCI to reach the targets. Transferability of acquired control strategy is essential for clinical applications outside of training settings and has been established as key learning criterion with healthy human participants (Sulzer et al., 2013b).

**Fig. 1.**
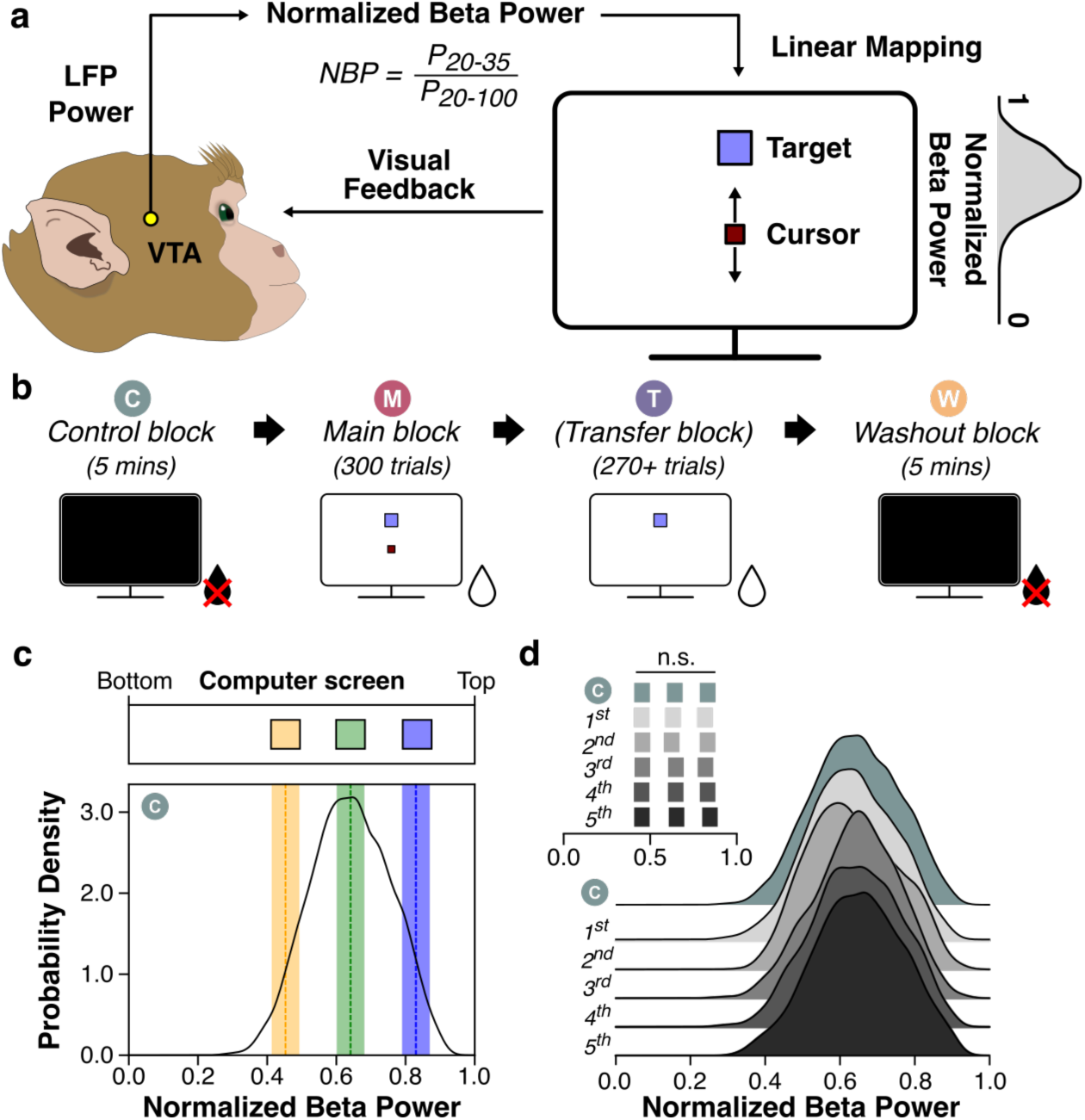
Experimental setup and normalized beta power (NBP). (**a**) Schematic of the neurofeedback paradigm. VTA LFP signals were used to calculate the instantaneous NBP, which were linearly mapped onto the computer screen as the vertical position of the cursor. (**b**) Experimental blocks. During the Control and Washout blocks, the computer screens and the juice reward system were turned off. The cursor provided visual feedback in the Main blocks. By contrast, in the Transfer blocks, only the target was visible. The blocks were color-coded throughout all figures. (**c**) Target locations. The probability density plot shows the NBP distribution from a Control block. The target sizes were the same and covered around 5-6 percentiles of data in High and Low targets. Target locations were also linearly mapped to the computer screen. The targets were color-coded throughout all figures. (**d**) Stability of NBP within a Control block. The five-minute recording of a Control block was segmented into five one-minute snippets. The NBP distributions and the three percentiles (inset) were estimated for each snippet. A Friedman test was used to test whether there are differences in percentiles across time (Q(4)=4.53, p=0.339).

Two male rhesus macaques (Monkey A: age 8, ∼13 kg; Monkey B: age 7, ∼10 kg) underwent training for the neurofeedback task. All procedures were approved by the Institutional Animal Care and Use Committee (IACUC) of The University of Texas at Austin. Non-human primates were used since they have similar cortical structures to humans and can perform complex tasks such as neurofeedback (Harding, 2017).

MR images for each subject were collected and then headpost implants were surgically placed to facilitate head fixation. Customized grids were designed based on the individual MR scans and a macaque brain atlas. For Monkey A, we used Linear Microelectrode Array V2 (LMAv2, MicroProbes for Life Science, Gaithersburg, MD), while V-probes (Plexon, Inc., Dallas, TX) were used for Monkey B. Electrodes were acutely lowered through the implanted chambers on one hemisphere of the brain to unilaterally target the VTA. The positions of the electrode tips were translated into the vertical distances required by each customized grid. Following electrode insertion, a 45-minute stabilization period without tasks served to settle the tissue. Training sessions typically lasted 2-4 hours.

Each training session was comprised of four blocks: Control, Main, Transfer and Washout. The Control and Washout blocks were designed to assess natural neural oscillations by examining the distribution of normalized beta power (NBP, details provided below). Moreover, the NBP distribution observed in the Control block determined the target locations for the Main block (see below). In the Main block, each trial commenced with the presentation of a target (task message: ‘start’), randomly selected from High, Center, or Low targets. Subjects were required to maintain the cursor within the target regions (‘hold’) for 200 ms to receive juice reward (‘reward’) while the target color changed from blue to green. Since there was no time limit for a trial, subjects could not avoid harder targets.

After completing 300 trials (100 for each target) in the Main block, subjects transitioned to the Transfer block, where the task structure remained identical, except the cursor was rendered invisible. While completion of the Transfer block was not mandatory for a session’s success, data from Transfer blocks were only included if subjects completed at least 270 trials. Regardless of successful completion of the Transfer block, each experimental session ended with the Washout block, which was identical to the Control block. Both blocks provided direct comparison of NBP distribution and other neural metrics before and after the neurofeedback training.

### Electrophysiology

Local field potentials (LFPs) were recorded from the ventral tegmental area (VTA) using 32-channel V-probes or LMAv2 electrodes alongside customized electrode grids. LFP signals were sampled at 1 kHz using Grapevine Analog and Digital I/O Modules (Ripple Neuro, Salt Lake City, UT). Raw LFP signals from each electrode were filtered using a low-pass filter at 250 Hz and a notch filter at 60/120/180 Hz to eliminate line noise. For each session, three channels within the VTA were manually selected for neurofeedback control, consistent with previous work (Khanna and Carmena, 2017). LFP power was computed for each channel using the multi-taper method provided by the *nitime* python package (with a window size of 200 ms, step size of 100 ms, and employing 6 tapers). The resulting LFP power time series were then averaged across the three selected channels (Khanna and Carmena, 2017). Since LFP is expected to be highly correlated with such spatially concentrated channels (Ahmadi et al., 2021), averaging across selected channels provides a better estimate of power and reduces noise.

In this study, the primary feedback signal was normalized beta power (NBP), calculated by dividing the power in the beta band (20-35 Hz) by the power in the broad band (20-100 Hz) for each time bin. In the Control block, the NBP time series exhibited a bell-shaped distribution with slight left-skewness. The 5^th^, 50^th^, and 95^th^ percentiles of this distribution were designated as the centers of the Low, Center, and High targets, respectively. High and Low targets, positioned 5 percentiles away from the extrema, were chosen to maintain comparable difficulty, with the Center target serving as a baseline. NBP was used rather than raw or z-scored beta power for multiple reasons. First, we rigorously assessed the three metrics based on their stability within a recording block and across recording sessions. To achieve this, we analyzed neural activity during Control blocks from Monkey A (n=15) and Monkey B (n=16). We identified the 5^th^, 50^th^ and 95^th^ percentiles of the distributions of these metrics. These descriptive statistics provide critical information about the distribution of each metric within the five-minute recording and are used to define the centers of targets in our neurofeedback experiments (Fig. 2). Across sessions, raw beta power exhibited day-to-day variability in power magnitudes, which was particularly evident in the 95^th^ percentiles of each session. While z-scoring slightly mitigated power magnitude discrepancies, the distribution of z-scored beta power was still as skewed as that of the raw beta power. This skewness posed challenges in ensuring uniform difficulty across targets and introduced potential bias in cursor placement due to the skewed distribution. On the other hand, NBP demonstrated stable percentiles across sessions, confined to a fixed interval regardless of power magnitude, and exhibited significantly reduced skewness. Since NBP is strictly confined between zero and one, it is straightforward and advantageous for mapping NBP to the computer screen across all sessions. We linearly map NBP to the vertical position of the computer screen so that the top of the screen corresponds to unity NBP while the bottom represents zero NBP.

**Fig. 2.**
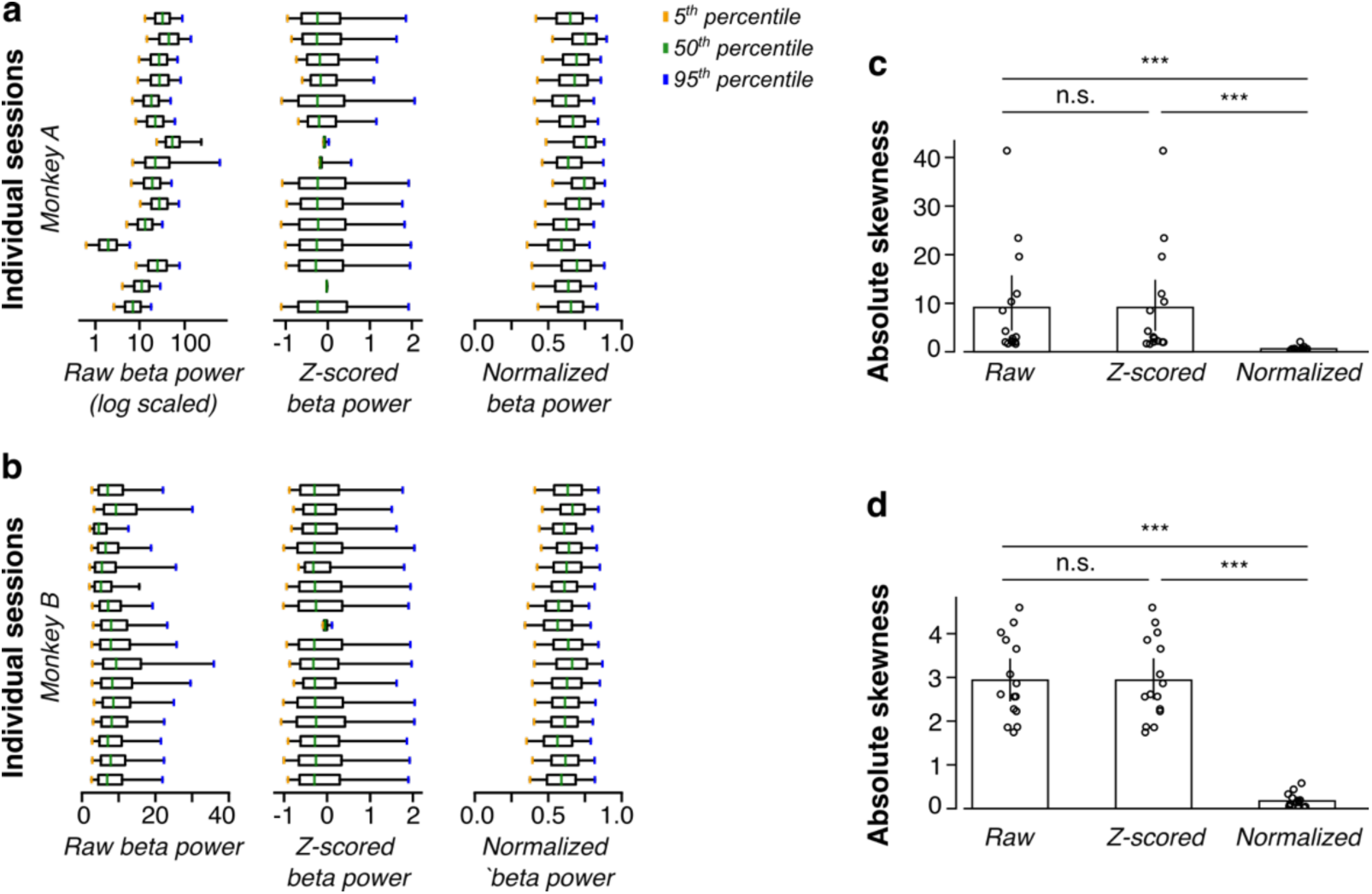
Normalized beta power is superior to raw or z-scored beta power. (**a** and **b**) The distributions of raw, z-scored, and normalized beta power in the Control blocks in each subject. Each box plot included the 5^th^, 50^th^, and 95^th^ percentiles of the distribution. Raw beta power suffered from day-to-day variance, while the z-scored beta power remained skewed (**c** and **d**). Absolute skewness considered left- and right-skew similarly. Normalized beta power was significantly less skewed than the other two metrics in both Monkey A (**c**) and Monkey B (**d**), as tested by one-way ANOVA. Monkey A: F(2,42)=4.274, p=0.020; post-hoc Tukey test: NBP and raw p=0.039, NBP and z-scored p=0.039, and raw and z-scored p=1.00. Monkey B: F(2,42)=64.96, p=1.40e-13; post-hoc Tukey test: NBP and raw p=4.93e-12, NBP and z-scored p=4.93e-12, and raw and z-scored p=1.00. Error bars represent standard errors of the mean.

Second, despite being normalized by a broader frequency range, NBP still strongly linearly correlated with beta power. To determine the frequency range for the normalization of beta power which preserved this linear relationship, we compared Spearman correlation of raw beta power with NBP computed with different denominator frequency ranges. Our result revealed peak correlation with a denominator range of 25-100 Hz that was consistent across subjects, indicating exclusion of power from lower frequencies enhanced correlation (Fig. 3a). This contrasts sharply with the weak correlation observed with a 4-59 Hz or a 17-47 Hz denominator frequency range (Fig. 3, b to d). The monotonic trend between raw beta power and NBP underscores subjects’ ability to increase (decrease) NBP by elevating (reducing) beta power. In addition, the weak correlation between NBP and gamma power, defined as power in the frequency range 35-100 Hz, indicated that the changes in NBP were mainly driven by changes in beta rather than gamma power (Fig. 3e).

**Fig. 3.**
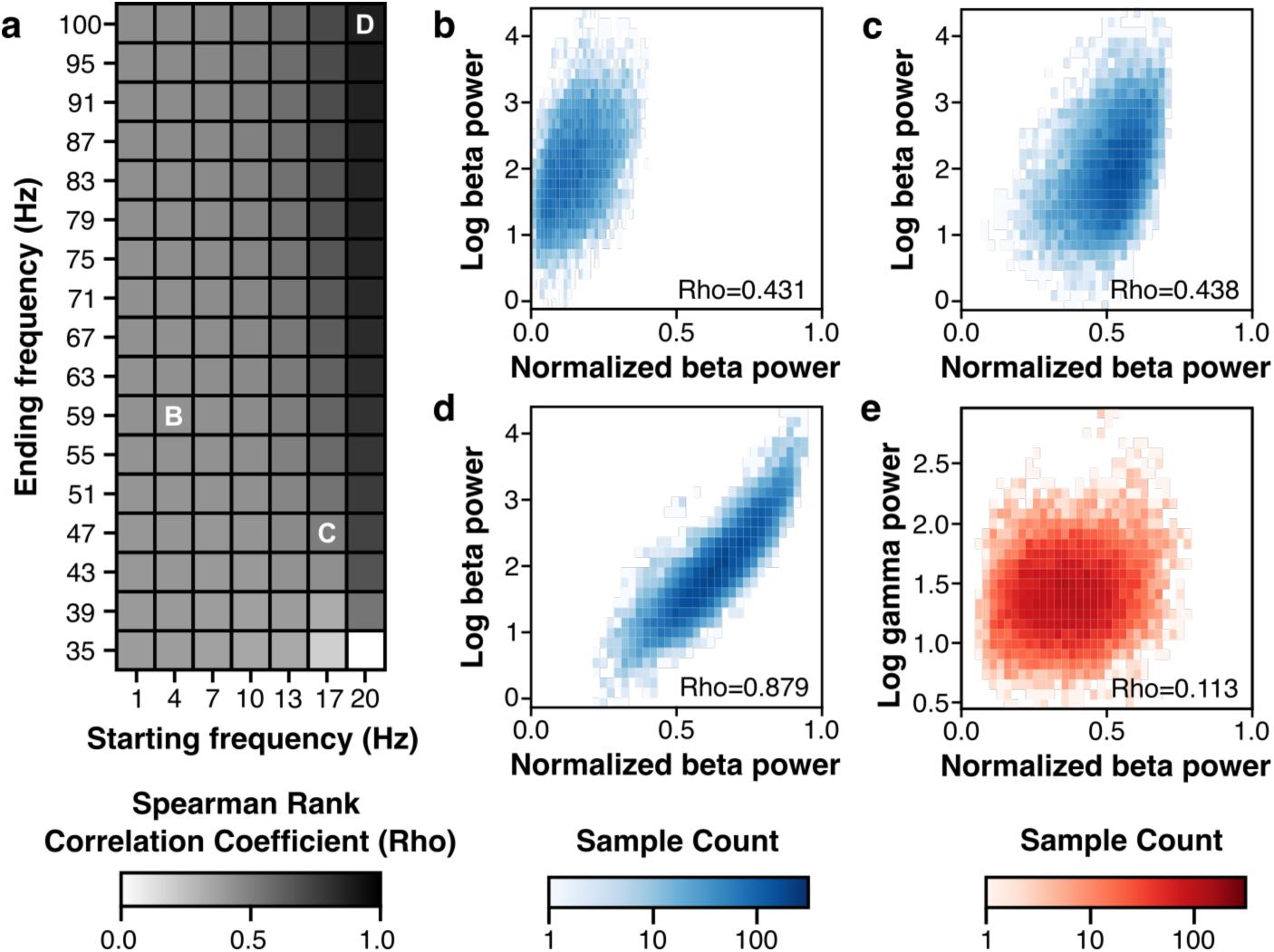
Optimizing NBP using rank correlations between NBP and beta power. (**a**) The Spearman rank correlation coefficients relating frequencies at start of trial (x-axis) and end of trial (y axis). The frequencies were selected so that the denominating frequency band for normalization was broader than the beta band. Thereby, we could ensure that the NBP was strictly bound between zero and one. (**b** to **d**) 2D histograms of the beta power (in log scale) and NBP using different representative denominating frequency bands, as indicated by the grids with corresponding letters in (**a**). (**e**) 2D histogram of the gamma power (in log scale) and the NBP using the optimal frequency range (**d**). In (**b** to **e**), the Spearman rank correlation coefficients are provided. The gradient colors indicate the number of samples accumulated in each pixel of the 2D histograms.

Kernel density estimate (KDE) plots were generated to visualize NBP distributions in a continuous fashion. The stability of NBP within Control blocks was assessed by dividing the five-minute recordings into five equally spaced segments, and KDE plots were constructed for each segment.

To quantify the changes of power in specific frequency bands during the ‘hold’ period, we first obtained the spectrogram aligned to ‘reward’ associated with the current target. The trial-averaged spectrogram was then normalized by the power spectral density in the Control block. The power during the ‘hold’ period was summed across frequency and averaged across time.

### Behavioral Metrics

The primary behavioral metric in this study is trial duration, defined as the time elapsed from the task message ‘start’ to ‘reward’. To visualize the trend of trial durations across trials, we applied a 20-sample boxcar filter to trial durations. Chance levels in Control and Washout blocks were estimated post hoc, by identifying instances where NBP remained within target regions for more than 200 ms in either block. Since there was no visual feedback (cursor), remaining in target regions for more than 200 ms was considered to occur purely by chance. The number of times the cursor remained in target regions for more than 200 ms was then divided by the length of those blocks (typically five minutes), quantifying chance level in rewards per minute (RPM). Similarly, RPM in Main and Transfer blocks were calculated by dividing the number of accomplished trials by the duration of the block.

Maximal improvement within a block was calculated in two steps. First, time-smoothed trial durations within a block were segmented into trial sets, with each set comprising ten samples for each target. Maximal improvement was defined as the difference between the mean trial duration in the first trial set and the set with the lowest trial duration, divided by the mean in the first set. If trial durations did not improve throughout the block, maximal improvement was reported as 0%.

### Statistical Analysis

Statistical analyses were performed using custom written Python3 scripts and a significance level set at 0.05. Effect sizes were computed using Cohen’s D (Cohen, 1988). P-values were denoted by asterisks, with one, two, and three asterisks representing p-value thresholds of 0.05, 0.01, and 0.001, respectively. One-tailed one-sample t-test, paired t-test, Friedman test, Pearson’s correlation coefficient test, and one-way ANOVA followed by post-hoc Tukey test were used for statistical comparisons. No statistical tests were run to predetermine the sample size.

### Code Accessibility

The code supporting the findings of this study are available from the corresponding author upon reasonable request.

## Results

### Normalized beta power

Central to our study is the development of the normalized beta power (NBP), specifically designed to stably quantify the modulation of VTA LFP signals. In contrast to conventional metrics such as raw (Engelhard et al., 2013) or z-scored band power (Chauvière and Singer, 2019), which can suffer from arbitrary thresholding (He et al., 2020) and inconsistent mappings across sessions, NBP provides a stable representation of brain activity that is strictly confined to a fixed range between zero and one (Fig. 2). NBP offers a robust and interpretable neural activity feature for regulation and facilitates the investigation of interactions between frequency bands, offering insights into the complex dynamics of brain self-regulation (see Materials and Methods). The target positions of each session were determined using the 5^th^, 50^th^, and 95^th^ percentiles of the NBP distribution from the Control block (Fig. 1c), labeled as the Low, Center, and High targets, respectively. NBP was intrinsically stable, especially when subjects were not engaging in experiments. By segmenting the five-minute NBP time series in a Control block into five equally spaced snippets, we found no significant difference in NBP among different times points across the 5^th^, 50^th^, and 95^th^ percentiles (Fig. 1d). This stationarity justifies our reliance on percentiles from the Control block for target position selection and underscores its potential as a valuable tool for elucidating neural mechanisms underlying self-regulation and guiding therapeutic interventions.

### Volitional regulation of VTA LFP improves with training

In the Main block, we used changes in the time the monkeys took to reach targets and thereby complete trials as measure of learning. These durations decreased across trials (Monkey A: n=15, Monkey B: n=16; Fig. 4a), with performance improvements significantly above chance levels (Fig. 5). By separating trial durations based on targets, we found distinct trends associated with each target that were shared across both subjects. While durations stayed relatively constant for Center targets, they decreased within a block for High and Low targets (Fig. 4a). Decreasing trial durations across trials indicate that subjects gradually improved their control over VTA LFP.

**Fig. 4.**
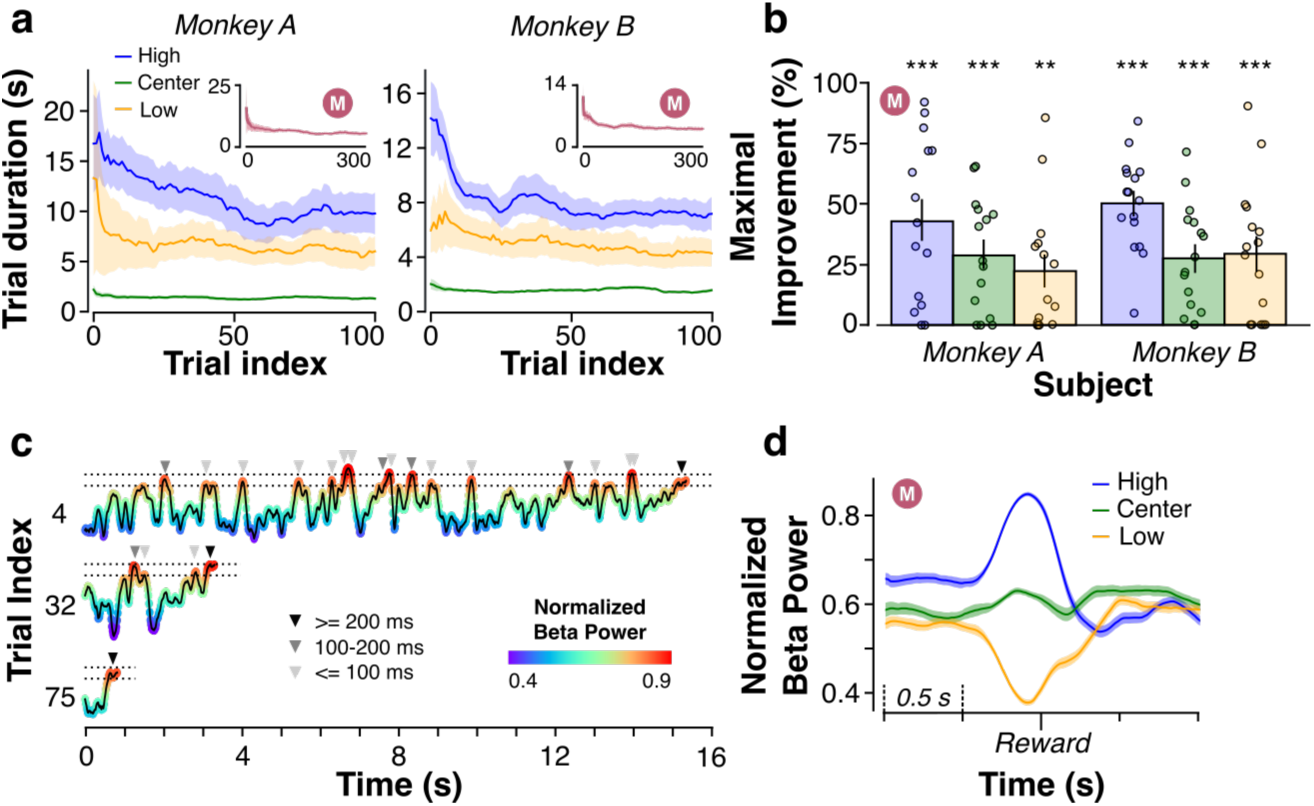
Successful volitional regulation and improvement of VTA NBP with training. (**a**) Trial durations in all Main blocks (Monkey A: n=15; Monkey B: n=16) were separated for each target and smoothed by a 20-sample boxcar filter. Insets show average trial durations without separating the targets, indicating general improvement over the Main blocks. (**b**) Maximal improvements for different targets across Main blocks. Bar plots represent the average. One-tailed one-sample t-test was used to verify if maximal improvements were greater than zero. Monkey A: High and Center: p<0.001, Low: p<0.01. Monkey B: p<0.001 for all targets. (**c**) Example trajectories of early, middle, and late trials for hitting High targets from a Main block, color-coded with NBP as shown by the color bar. Dashed lines represent borders of the target (here NBP=0.800 to 0.884). Arrows above the trajectories mark the periods when the cursor was in the target region and shade reflects duration within the target region. (**d**) NBP aligned to reward (end of trials) and averaged across trials for individual targets. Each interval in the x axis represents 0.5 sec. Shaded areas and error bars are standard error of the mean.

**Fig. 5.**
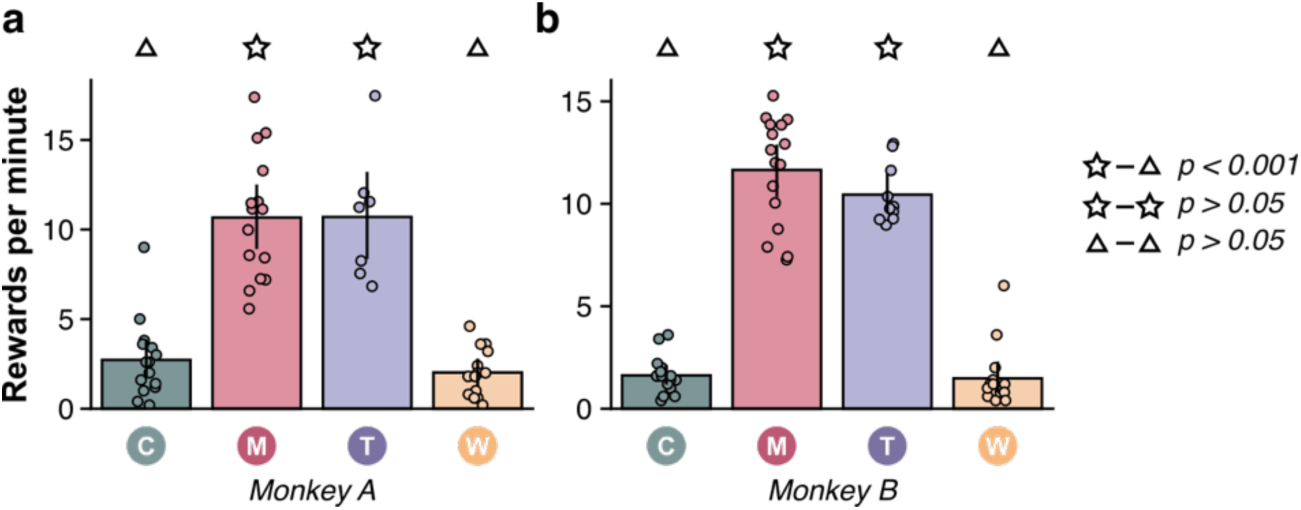
Rewards per minute in each block. The rewards per minute (RPM) in the Control and Washout blocks correspond to chance and were computed post-hoc because the subjects did not actually receive any rewards during these blocks. We first calculated the target locations using the three percentiles in the Control block, and then examined through the NBP time series to count the number of times the cursor stayed within the defined target regions for more than 200 ms. The RPMs in the Main and Transfer blocks were calculated by the actual number of rewards received divided by the duration of the block. (**a**) Monkey A: F(3,47)=39.57, p=6.54e-13; post-hoc Tukey tests: Control vs. Main p<0.001, Control vs. Transfer p<0.001, Control vs. Washout p=0.89, and Main vs. Transfer p=0.99. (**b**) Monkey B: F(3,53)=140.26, p=3.43e-25; post-hoc Tukey tests: Control vs. Main p<0.001, Control vs. Transfer p<0.001, Control vs. Washout p=0.996, and Main vs. Transfer p=0.340. Error bars represent standard errors of the mean.

To assess the degree of reduction in trial durations, we quantified “maximal improvement” for each target within each block (see Materials and Methods). Our analysis revealed significant improvements across the Main blocks for each target location in both subjects (Fig. 4b). These findings indicate that the observed modulation of LFP power is not random, but rather an intentional strategy employed by the subjects to optimize performance.

Beyond confirming behavioral improvements, we delved deeper into NBP trajectories from three representative trials within a single Main block: one from early, one from the middle, and one from late in the block (Fig. 4c). Visual inspection revealed not only faster trial completion but also remarkable NBP control. Shorter NBP trajectories across trials indicated that the monkeys needed fewer attempts as they became more practiced. Note that while regulation itself was important, the task demanded precise and sustained control of NBP to certain intervals for at least 200 ms. To capture this temporal aspect, we averaged NBP around the time of ‘reward’ (trial completion) for each target (Fig. 4d). On average, NBP regulation typically initiated 0.3-0.5 seconds before trial completion and returned to baseline within a similar period afterwards. This pattern suggests that the subjects learned to quickly and intentionally adjust NBP to specific ranges, demonstrating that monkeys can volitionally regulate NBP in the VTA. The timescale is also consistent with beta bursts, which are functionally related to aspects of cognition (Lundqvist et al., 2024), such as movement planning (Little et al., 2019) and working memory (Miller et al., 2018), providing further evidence of physiological relevance of the neurofeedback signal used.

### Differential contributions of beta and gamma power to NBP regulation

We optimized the NBP to achieve a high correlation with beta power (Fig. 3). However, NBP is a function of both beta (20-35 Hz) and gamma (35-100 Hz) band power. Studies have shown that gamma power in the VTA remains relatively stable during decision-making tasks compared to other frequency bands (Oberto et al., 2023). Here, we investigated whether the subjects primarily regulated beta power to influence NBP or if alterations in both beta and gamma bands contributed differentially to changes in NBP.

To explore the contribution of each frequency band to the regulation of NBP, we synchronized and normalized spectrograms with the time of reward for each target in the Main blocks (Fig. 6a). As expected, based on the correlation between beta power and NBP, beta power increased for High targets, decreased for Low targets, and remained stable for Center targets compared to Control block levels (Fig. 6b). Next, we assessed the potential involvement of the gamma band. Even though gamma power was only weakly correlated to NBP (Fig. 3e), it significantly contributed to changes in NBP. Compared to beta power, gamma power exhibited a less symmetrical pattern across targets (Fig. 6b). Specifically, it increased for Low and Center targets but remained stable for High targets. Notably, this modulation was specific to the beta and gamma bands, as delta and theta did not show significant changes, and while alpha showed significant changes, its modulation was much weaker (Fig. 7a). Thus, the subjects flexibly, differentially, and exclusively controlled beta and gamma frequency bands to accomplish the task.

**Fig. 6.**
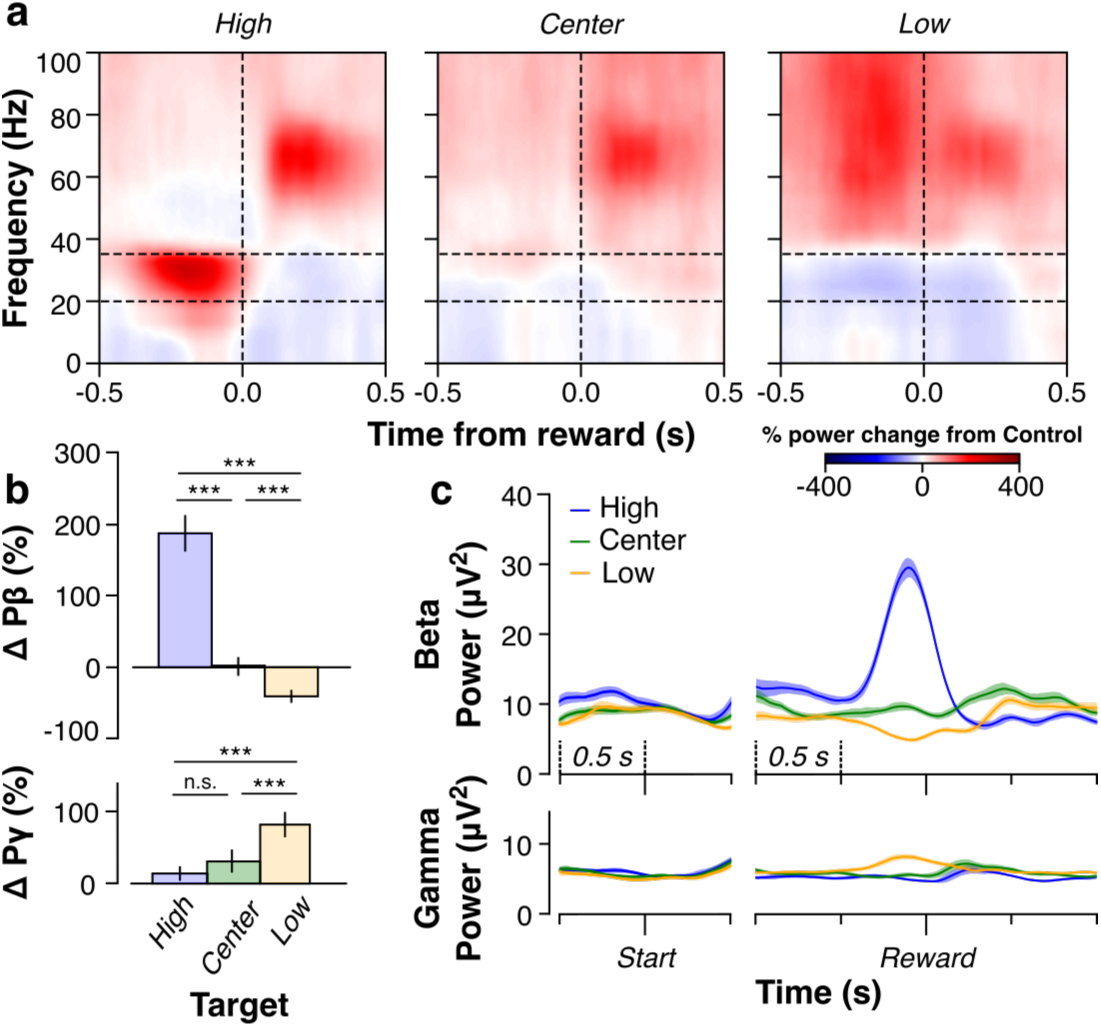
Differential contribution of frequency bands to volitional VTA regulation. (**a**) Spectrograms were aligned to reward for each target and normalized by the power spectral density in the Control block to show relative changes in power for each frequency bin. Dashed lines represent the boundaries of the beta power band (20 and 35 Hz). (**b**) Changes in beta and gamma power were summed across the beta band (20-35 Hz) and averaged across trials (n=100 trials). Error bars are standard error of the mean. One-way ANOVA was used to test whether changes in power differed across targets. Beta power: F(2,298)=296.52, p=1.32e-71, post-hoc Tukey test p<0.001 for all combinations. Gamma power: F(2,298)=43.60, p=2.45e-17, post-hoc Tukey test p<0.001 for Low, but not for High and Center (n.s.). (**c**) The raw beta and gamma powers were aligned and averaged to start of trial and reward (end of trial). Shaded areas are standard error of the mean.

**Fig. 7.**
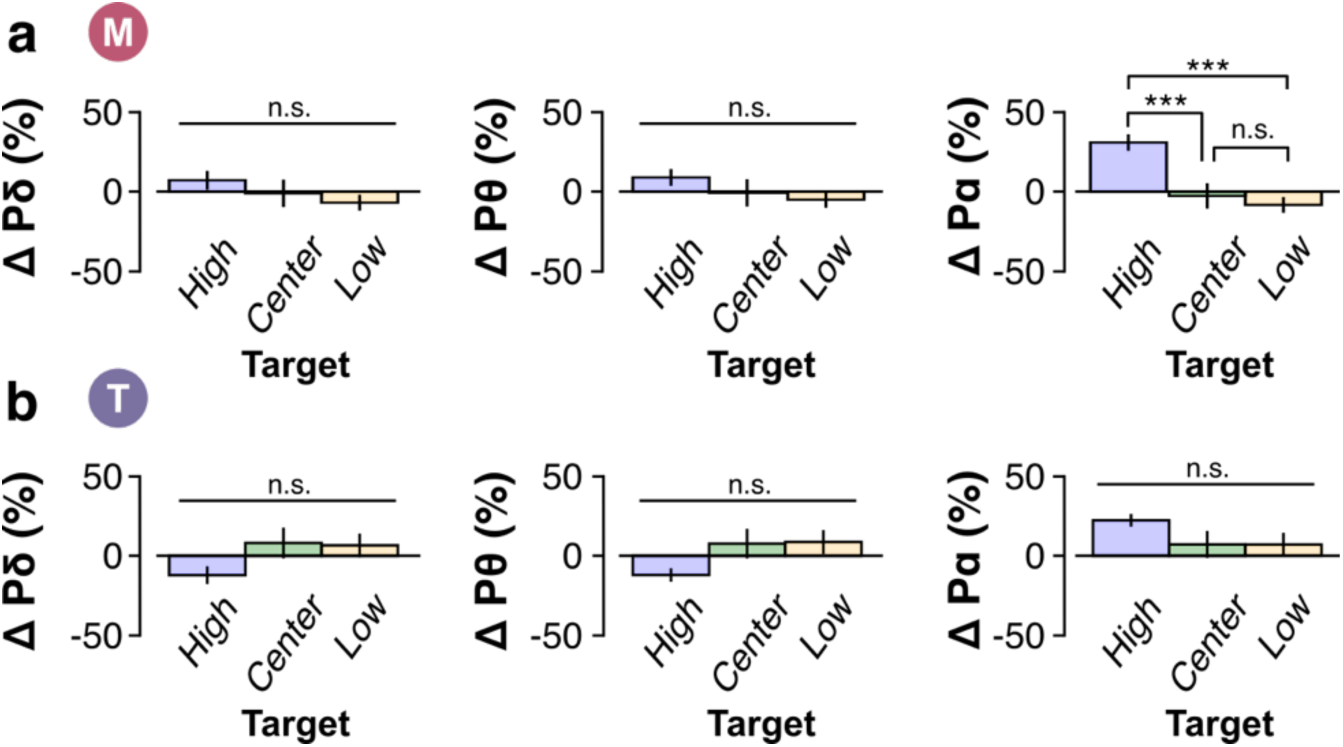
Stability of powers other than beta and gamma frequency bands during regulation. (**a**) Similar to Fig. 3B, changes in each band power in the Main block were summed across each band (left: delta 1-4 Hz, center: theta: 4-8 Hz, and right: alpha: 8-12 Hz) and averaged across trials (n=100 trials). Error bars are standard error of the mean. One-way ANOVA was used to test whether changes in power differed across targets. Delta power: F(2,298)=1.11, p=0.33. Theta power: F(2,298)=1.20, p=0.30. Alpha power: F(2,298)=11.66, p=1.3e-5, post-hoc Tukey test p<0.001 for High, but not for Low and Center (n.s.). (**b**) Similar to Fig. 4E, changes in power in the Transfer block were summed and averaged across trials. Error bars are standard error of the mean. One-way ANOVA was used to test whether changes in power differed across targets. Delta power: F(2,297)=2.54, p=0.08. Theta power: F(2,297)=2.12, p=0.12. Alpha power: F(2,297)=1.63, p=0.20.

In addition to examining overall power changes relative to the Control block, we investigated the temporal dynamics of regulation for both the beta and gamma frequency bands. We summed the power in each frequency band, synchronized them to both the ‘start’ and ‘reward’ times, and averaged them across trials. We observed indistinguishable power differences across targets at the beginning of trials (‘start’) for both bands. However, both frequency bands began to be regulated approximately 0.3-0.5 seconds before trial completion (Fig. 6c). This temporal pattern is compatible with intentional power regulation. At the beginning of a trial, subjects did not intentionally regulate power in either band to specific levels but did so as they approached trial completion. Band power returned to baseline levels upon trial completion in both cases.

We noted that raw beta power was stronger than raw gamma powers, consistent with the 1/f trend seen in neural signals where higher frequency bands often exhibit relatively weaker expression compared to lower frequency bands (Novikov et al., 1997). This discrepancy was more pronounced during intentional modulation periods around the time of reward. Beta power could increase by up to 200% of its original magnitude, whereas gamma power saw a maximum increase of around 100%. The inability to decrease power for High targets may reflect the fact that gamma power was already relatively low. Instead, subjects predominantly increased beta power to achieve High targets.

### Successful regulation of VTA LFP without continuous feedback

Transfer blocks immediately followed the successful completion of Main blocks. In lieu of verbally instructing participants to regulate (as in human neurofeedback experiments), in the Transfer blocks we visually presented only the target (as in the Main block) while occluding the cursor position. Accordingly, the subjects performed the task without continuous visual feedback in the Transfer blocks.

At the beginning of Transfer blocks, trial durations for all targets were longer than at the end of Main blocks (Fig. 8a). However, the trial durations decreased throughout the block, indicating that the regulation strategy developed with continuous feedback can generalize and be refined over time even without visual feedback. Maximal improvement analyses in Transfer blocks (Fig. 8b) also revealed significant improvement in trial durations for both subjects.

**Fig. 8.**
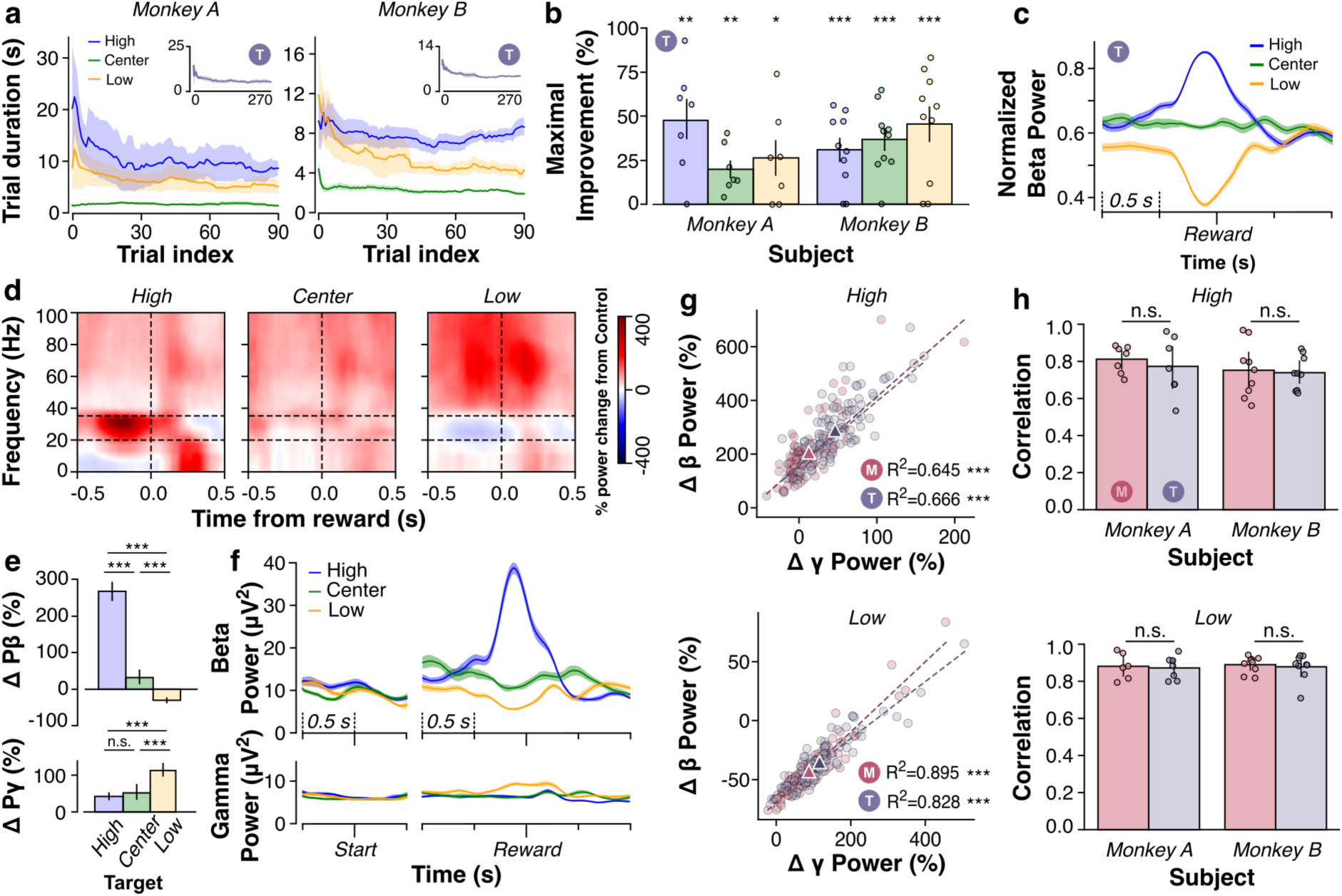
Regulation transfers to occluded cursor condition and improves over time. (**a**) Trial durations (Monkey A: n=7; Monkey B: n=10). (**b**) Maximal improvements for different targets. One-tailed t-tests assessed the difference from zero. Monkey A: High: p=2.82e-3, Center: p=3.74e-3, Low: p=0.02; n=7. Monkey B: High: p=5.37e-4, Center: p=9.8e-5, Low: p=6.09e-4; n=10. (**c**) NBP aligned to reward and trial-averaged for individual targets. (**d** to **f**) Spectral analyses from representative Transfer block. (**e**) One-way ANOVA tested differences in changes in power across targets. Beta power: F(2,297)=373.65, p=8.11e-82, post-hoc Tukey test p<0.001 for all combinations. Gamma power: F(2,297)=29.12, p=2.83e-12, post-hoc Tukey test p<0.001 except High and Center (n.s.). (**g**) Co-modulation patterns are preserved between Main and Transfer blocks (data from representative session). Changes in power are relative to Control block. Triangles represent centroids of each group. High target: slope is 2.76 (R^2^=0.645, p=8.67e-24) and 2.62 (R^2^=0.666, p=4.68e-25) for Main and Transfer blocks. Low target: slope is 0.3 (R^2^=0.895, p=2.92e-50) and 0.24 (R^2^=0.828, p=3.24e-39) for Main and Transfer blocks. (**h**) Correlation between changes in beta and gamma power. Paired t-tests reveal no difference between Main and Transfer. Monkey A: High p=0.434, Low p=0.424 (n=7). Monkey B: High p=0.786, Low p=0.698 (n=10). Error bars represent SEM.

In addition, we synchronized and averaged the NBP around the time of reward for each target (Fig. 8c). Notably, both monkeys precisely regulated NBP to the designated targets also in the Transfer blocks as they had in the Main blocks. Mirroring the behavior observed in the Main blocks, regulation typically commenced around 0.3-0.5 seconds before trial completion. Furthermore, NBP reverted to baseline levels upon trial conclusion, compatible with intentional regulation of brain activity. This pattern suggests that subjects deliberately adjusted their brain activity also in the Transfer blocks, with no further intentional changes until the subsequent trial.

The synchronized spectrogram analyses of Transfer block neural data yielded analogous patterns to those observed in the Main blocks prior to target attainment (Fig. 8d). Upon aggregating power across frequency bands within the hold-target time window (t=-0.2 to t=0 seconds), discernible alterations emerged across various targets in beta and gamma power (Fig. 8e) but not in other frequency bands (Fig. 7b). Temporally, regulation again occurred about 0.3-0.5 seconds before trial completion and reverted to baseline levels post-trial (Fig. 8f). Thus, as in the Main blocks, beta and gamma bands were differentially regulated.

However, a significant discrepancy emerged between Main and Transfer blocks. In Transfer blocks, a pronounced increase in power within the lower frequency range, particularly the theta band (4-8 Hz), around 200 ms after trial completion was notable (Fig. 8d and Fig. 9a). This theta power increase was particularly accentuated for High targets but also discernible for Center and Low targets. Since theta band power in the VTA has been linked to reward-prediction error (RPE) (Kim et al., 2012), we suggest that the absence of the cursor might have led to an increase in RPE magnitude in Transfer blocks, with the unexpected change in target color serving as the only visual indicator of trial completion. Analysis of theta band power across trials provided several potential insights (Fig. 9a). First, theta band power tended to remain low across all targets in the Main blocks, which might indicate stronger RPEs in Transfer blocks due to the absence of the cursor. Second, theta power in Transfer blocks appeared notably high at the beginning, especially for High and Low targets, reflecting higher uncertainty when regulating without visual feedback, but it generally decreased over time while remaining higher than in Main blocks (Fig. 9, b to c). Third, theta power in Transfer blocks seemed to activate only when monkeys successfully held regulation, suggesting reduced reward expectations and increased positive RPEs when trials ended, and reduced negative RPEs when they did not (Fig. 9d).

**Fig. 9.**
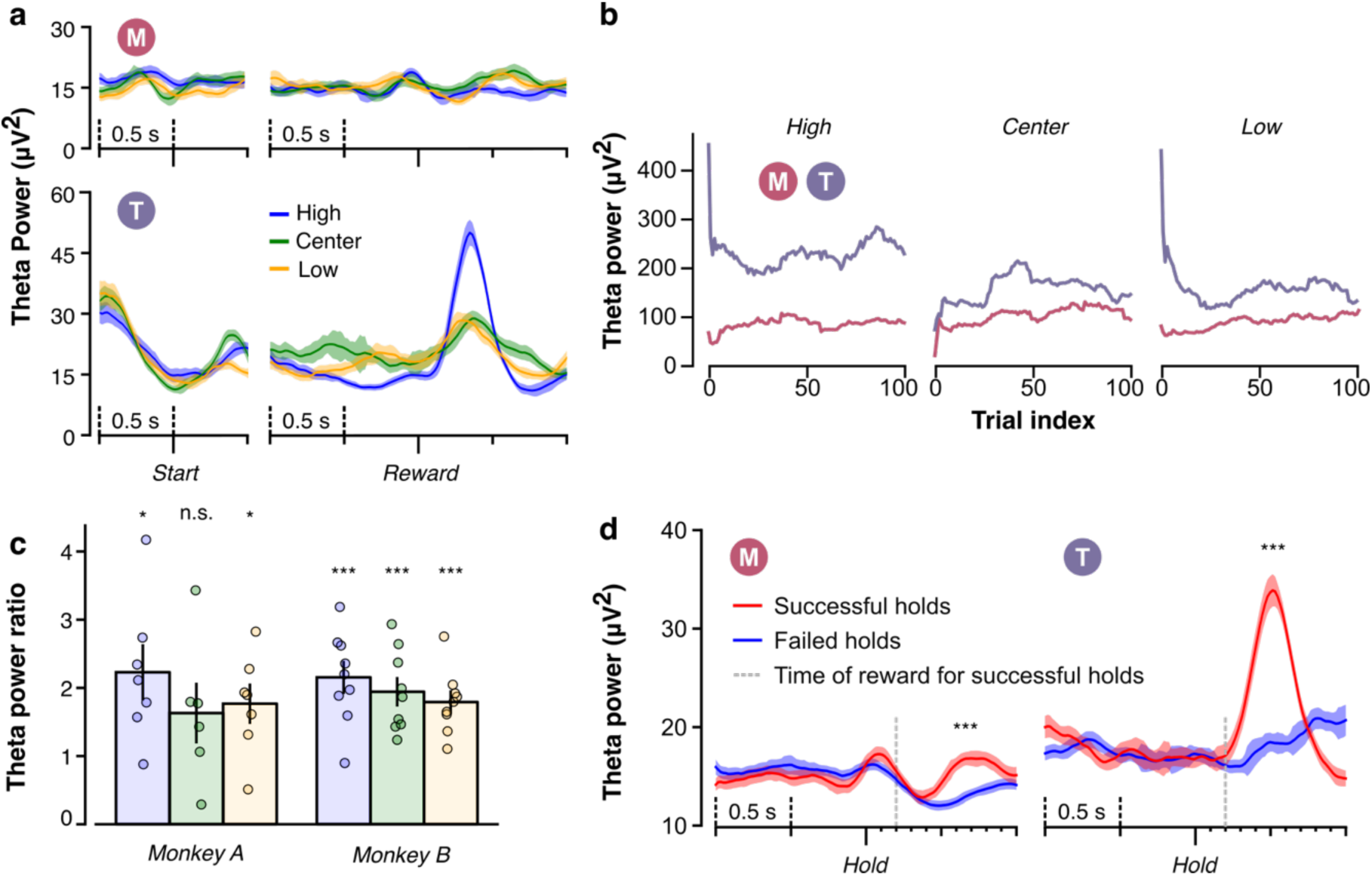
Theta power in the Transfer blocks shows positive RPE after reward. (**a**) The raw theta power was aligned and averaged to start of trial and reward (end of trial) in both Main (top) and Transfer (bottom) blocks, similar to the plotting scheme in Fig. 3C. Shaded areas are standard error of the mean. (**b**) Theta power accumulated between t=0.2 to t=0.5 sec from Reward for each target condition in the Main and Transfer blocks of a representative session and smoothed using a 20-sample boxcar filter. This interval was selected due to its relevance to the RPE signal (*48*). (**c**) Average ratio of theta power between Transfer and Main blocks. One-tailed one-sample t-tests assessed whether the ratios were greater than one. Monkey A: High target p=0.010, Center target p=0.100, and Low target p=0.016. Monkey B: High target p=4.32e-4, Center target p=7.12e-4, and Low target p=4.59e-4. Error bars represent standard errors of the mean. (**d**) Theta power was aligned to Hold and averaged based on whether the holds are successful. If successful, rewards were delivered at t=0.2 sec. Theta power between successful and failed holds from t=0.4 to t=0.7 sec (identical to t=0.2 to t=0.5 sec from Reward, as shown in **b**) were statistically different in both Main and Transfer block based on one-sample t-tests. Main p=4.90e-8 (Cohen’s D = 0.415); Transfer p=1.754e-12 (Cohen’s D = 0.486).

We then investigated whether the subjects applied the strategies they acquired in the Main blocks also in the Transfer blocks. We averaged beta and gamma power for each trial in Main and Transfer blocks for High and Low targets (Fig. 8g). The centroids (triangles in Fig. 8g) of each block aligned with the average power in previous results (Fig. 6b and 8e). Significant linear relations between changes in beta power and changes in gamma power were observed for both blocks for both targets, indicating co-regulation of beta and gamma power. Importantly, the distributions of changes in beta and gamma power were preserved from Main to Transfer blocks. The similar co-regulation patterns suggest successful transfer of volitional NBP regulation from the Main block to the subsequent Transfer block. We also assessed the correlation between changes in beta and gamma powers and found no differences between Main and Transfer blocks within subjects for both targets (Fig. 8h). This finding demonstrated that the subjects could not only learn to volitionally control their LFP signals in the VTA but also transfer these skills to a similar yet more challenging context.

### LFP activity reverts to initial dynamics

At the end of each session, a five-minute Washout block again assessed natural brain fluctuations. To evaluate the impact of neurofeedback training on neural activity, we examined changes in power within the beta and gamma bands between the Washout and Control blocks (Fig. 10). Our results provide no evidence for a change in power within these bands. Interestingly, while there was an overall increase in power, the effect size in NBP was comparatively smaller, underscoring the remarkable stability of this metric. Analysis of NBP distributions across the four blocks revealed consistent percentiles (Fig. 11a). Over all sessions, variance was similar for Control and Washout blocks but tended to be higher for Main and Transfer blocks (Fig. 11b). Thus, regulation of NBP was associated with increased NBP variance during neurofeedback blocks, with variance returning to pre-training levels once the subjects stopped regulation.

**Fig. 10.**
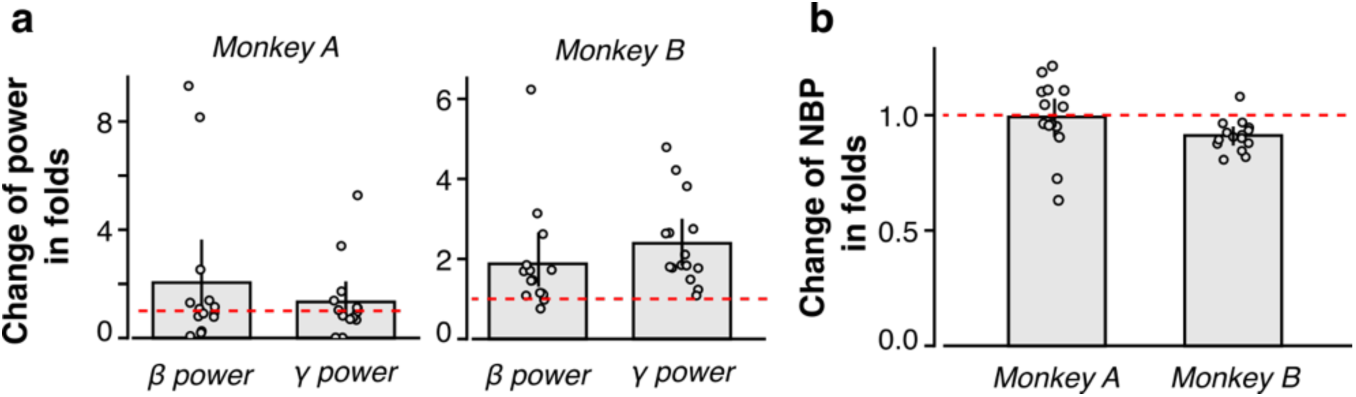
Changes in beta and gamma power and NBP after neurofeedback training. (**a**) For each session, the average beta and gamma power in the Washout block was divided by the average power in the same frequency band in the Control block. (**b**) The mean NBP in the Washout block was divided by the mean NBP in the Control block for each session. One-tailed one-sample t-test served to test whether the changes were greater than one: Monkey A: beta power p=0.097 (Cohen’s D = 0.365), gamma power p=0.180 (Cohen’s D = 0.254), and NBP p=0.642 (Cohen’s D = 0.100), n=14; Monkey B: beta power p=0.013 (Cohen’s D = 0.640), gamma power p=1.28e-4 (Cohen’s D = 1.253), and NBP p=0.999 (Cohen’s D = 1.335). Error bars represent standard errors of the mean.

**Fig. 11.**
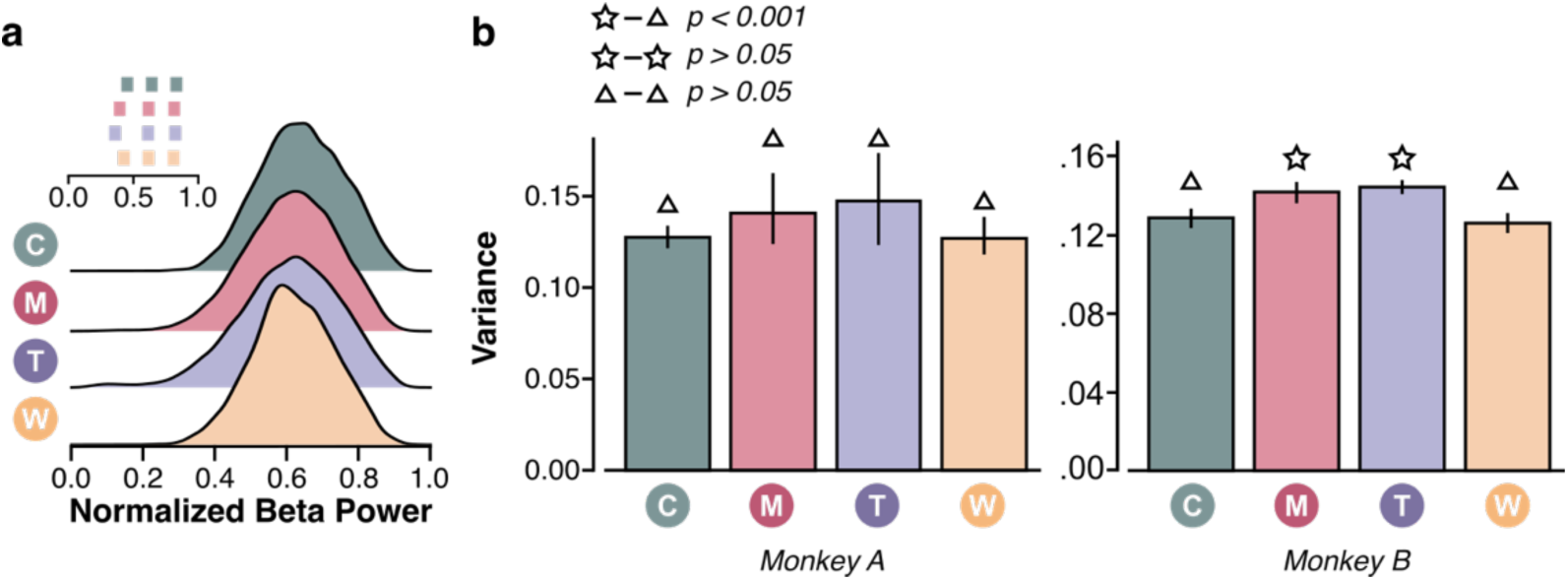
Distribution and variance of NBP in the four blocks. (**a)** NBP distributions in a representative session. Inset shows the 5^th^, 50^th^, and 95^th^ percentiles of each distribution. (**b**) Variance of each block in all sessions. One-way ANOVA was used to test for differences across blocks. The code above each bar indicates significance levels using post-hoc Tukey tests. Monkey A: F(3,47)=1.57, p=0.21. Monkey B: F(3,53) = 16.65, p=9.6e-8; post-hoc Tukey tests: Control vs. Main p=2.05e-4, Control vs. Transfer p=1.23e-4, Control vs. Washout p=0.92, Main vs. Transfer p=0.89, Main vs. Washout p=1.3e-5, and Transfer vs. Washout p=1.0e-5. Error bars represented standard errors of the mean.

## Discussion

Our study introduces several significant contributions to the field of neurofeedback. First, we developed a customized control metric characterized by consistent mapping and inherent stability. Second, we successfully demonstrated the feasibility of VTA LFP neurofeedback in non-human primates, a novel achievement in this research area. Third, we conducted what we believe is the first non-human neurofeedback experiment that includes a transfer phase, showcasing the ability of subjects to apply learned strategies in subsequent sessions. Fourth, we showed that the acquired control strategy could be effectively transferred to subsequent blocks, indicating the potential for long-term retention and application of neurofeedback training. Collectively, our findings leverage the LFP neurofeedback paradigm to explore the flexibility and controllability of VTA LFP power, underscoring its potential as a therapeutic target for dopamine-associated neuropsychiatric disorders.

Unlike traditional neurofeedback experiments that typically focus on increasing power (Engelhard et al., 2013; Chauvière and Singer, 2019), our experiment design involved increasing and decreasing NBP and revealed distinct strategies for different targets. Our results demonstrate that monkeys can volitionally and bi-directionally regulated beta activity in order to reach the High and Low targets but additionally increased gamma power to reach the Low targets. This finding sheds light on the intricate dynamics of frequency band interactions, suggesting that subjects adopt nuanced strategies to optimize control over specific power bands. Despite their lack of explicit knowledge regarding the frequency band mapping, they demonstrated a remarkable ability to fine-tune beta and gamma power regulation to attain desired NBP levels. Importantly, our results align with prior VTA neurofeedback studies employing fMRI, indicating that primates can voluntarily regulate signals originating from dopaminergic regions and transfer these effects post-training (Sulzer et al., 2013b; MacInnes et al., 2016).

The LFP neurofeedback paradigm facilitates investigation of power features that are not directly associated with the feedback signal (Khanna and Carmena, 2017; Chauvière and Singer, 2019), in addition to validating intentional control of brain signals. In our investigation of learning transfer, we are particularly interested in the theta power in the VTA due to its connection with RPE (Kim et al., 2012). Even though the subjects applied similar regulation strategies to the Transfer blocks, they were not fully aware of the instantaneous cursor location which visually represented NBP during training. This increased uncertainty led to an increase in the theta power after being rewarded. In line with learning reducing uncertainty (Soltani and Izquierdo, 2019), the theta power decreased as the subjects practiced more within the Transfer block. This integrated information broadens our understanding of VTA neurofeedback, especially for the learning process in the Transfer blocks.

In Transfer blocks, the uncertainty of brain states potentially reflects individual variability in response to neurofeedback. Some individuals may not respond to neurofeedback interventions as expected, a phenomenon referred to as non-responder effects (Alkoby et al., 2018). Even though we demonstrate differential contributions between frequency bands and transferable patterns across blocks on average, we also found variability in performance between sessions that cannot be explained by features of the oscillatory neural activity. These observations suggest that understanding the limits of neural modulation is crucial, as it may influence or cause the occurrence of non-responder effects in neurofeedback.

## Acknowledgments

The authors acknowledge C. Jacobs, L. A. Wilding, and B. Shukla for surgical support. S.R.S. was supported through the Whitehall Foundation (2022-12-071) and National Science Foundation CAREER Award (2145412); P.N.T. was supported through Swiss National Science Foundation (165884; 176016; 188878; and 207613).

